# Peer teaching as bioinformatics training strategy: incentives, challenges, and benefits

**DOI:** 10.1101/2021.08.26.457832

**Authors:** Nur-Taz Rahman, Caitlin Meyer, Durga Thakral, Wesley L. Cai, Ann T. Chen, Razib Obaid, Rolando Garcia-Milian

**Affiliations:** Bioinformatics Support Program, Research and Education Services, Harvey Cushing/John Hay Whitney Medical Library, Yale University; Dept. of Medicine, Brigham and Women’s Hospital, Harvard Medical School, Boston, MA 02115; Dept. of Pathology, Yale School of Medicine, New Haven CT 06510; Dept. of Biomedical Engineering, Yale University, New Haven CT 06510; RARAF Radiological Research Accelerator Facility, Nevis Laboratory, Columbia University, Irvington, NY 10533

**Keywords:** bioinformatics, peer teaching, training, data, education, computational biology

## Abstract

As biomedical research becomes more data-intensive, bioinformatics is becoming essential to understanding biological processes, systems, and diseases. In this paper we describe the use of a series of peer teaching workshops as a strategy to respond to the bioinformatics training needs at a research-intensive institution. In addition to the data collected from the workshops, we also used personal experiences of researchers who participated as peer teachers to understand the incentives, challenges, and benefits of peer teaching. Developing communication skills such as confidence in teaching, explaining complex concepts, and better understanding of the topic emerged as primary benefits that the teachers obtained from this experience. Lack of time for teaching and the struggles of classroom management were identified as two major challenges. We suggest that peer teaching can be beneficial not only to train researchers in bioinformatics, but also as a professional development opportunity for graduate students and postdoctoral trainees.

## Background

Twenty-first century biomedical sciences have transformed into data-intensive fields, generating multiple kinds of big data. Hence, biomedical researchers are becoming more dependent on computation-intensive bioinformatics approaches to answer research questions. Bioinformatics training is challenging not only because of its interdisciplinary nature, but also because of the recent and extensive technological changes. As bioinformatics becomes increasingly central to research, there is a pressing need to train non-bioinformaticians to learn some aspects of data analysis (Schneider et al., 2010).

Medical Libraries have been providing bioinformatics services in support of biomedical research for several years. These programs usually include consultation services, training, collaborations, and networked biological information resources (Chattopadhyay et al., 2006; Li, Chen & Clintworth, 2013; Hosburgh, 2018). In 2014, the Harvey Cushing/John Hay Whitney Medical Library (CWML) at Yale University started a Bioinformatics Support Program (BSP) intended to support Yale University-affiliated biomedical researchers. A study conducted by this program revealed researchers had unmet bioinformatics training needs (Garcia-Milian et al., 2018). Peer teaching was one of the primary strategies considered to fill this instructional gap. Biomedical researchers at any career level were invited to share their knowledge, from an altruistic viewpoint. Our peer teachers shared their knowledge with the community through a series of workshops that we called peer-to-peer (P2P) teaching.

P2P teaching (different to peer instruction) occurs when individuals, who are not professional teachers, help others from similar social groups with learning a particular topic (Topping, 1996). Biomedical researchers seeking bioinformatics expertise are spread out across multiple departments, making it difficult to find experts or faculty willing to provide bioinformatics training beyond the formal curriculum (Zatz, 2002). Junior researchers (graduate students and postdoctoral trainees) are much more fluid in terms of their departmental affiliation. In addition, advocates of P2P teaching argue that peer teachers share similar knowledge, challenges, and language (known as cognitive congruence), which has a beneficial effect on learning outcomes (Yu et al., 2011). A peer teacher providing bioinformatics training is more likely to be aware of the current challenges faced by those who are trying to learn the skill. Therefore, s/he is better able to tailor the instruction material to the needs of the learners. Some studies have found that teachers with specific short-course experience may be more suitable for bioinformatics training, compared to university professors (Via et al., 2013). A more seasoned bioinformatician may be harder for the learners to follow and keep pace with.

Previous studies have also indicated that P2P teaching achieves learning outcomes that are comparable to those produced by faculty-based teaching (Yu et al., 2011). We would like to note that P2P teaching should not be confused with peer instruction, which occurs in the form of student interaction (usually to promote problem-solving or discussion), within a modified traditional lecture format, where a non-peer teacher is still present (Mazur, 1997). Peer instruction has been previously used in bioinformatics, for example, peer-assisted learning as part of a research-based laboratory course (Shapiro et al., 2013), and concerted teaching of biology and computer science courses (Goodman and Dekhtyar, 2014). However, we did not find a previous report of using P2P teaching in bioinformatics instruction. In this paper we share our experience of using a P2P teaching strategy to support the training needs of biomedical researchers, at a large research-intensive academic institution. We also explore the incentives, challenges, and benefits of peer teaching from the teachers’ point of view.

## Methods

Instructors were recruited via personal requests for volunteers to teach single sessions within the framework of the workshops regularly offered by the Bioinformatics Support Program of the Harvey Cushing/John Hay Whitney Medical Library (CWML). Volunteers decided on the best format for presenting and teaching the content. In addition to data collection on the topics and attendees of the workshops, we used researchers’ personal experience (Byczkowska-Owczarek, 2014) as a method to gain more insight into the experience from those who participated as instructors. To this end, authors of this paper who participated as bioinformatics peer teachers (NR, DT, WLC, ATC, RO) were given a questionnaire with open questions to reflect on the benefits, challenges, and rewards of P2P teaching. Their responses were analyzed (CM, RGM, NR) to find common and unique topics. At the time of providing instruction all teachers were graduate PhD students. Access to the questionnaire used in this study can be found at the EliScholar repository (http://works.bepress.com/rolando_garciamilian/38/), a digital platform for scholarly publishing provided by Yale University Library.

Data regarding positions and departments of the registrants were extracted from the registration database where registrants were de-identified, and only the metadata was conserved. These data were summarized and used to create table 1, and figures 1 and 2. All figures were generated in R (version 3.6.2) using ggplot2 (version 3.3.1).

**Table 1.** Summary of sessions offered including description and number of registrants by session.

| Session title | Session type | Description | Registrants |
| --- | --- | --- | --- |
| Next Gen Sequencing for Biologist Series: Introduction to Exome Sequencing Analysis | Presentation | An overview on Whole Exome Sequencing analysis including workflow, file types and analysis tools | 70 |
| From benchside to Rstudio: Part 1 | Hands-on | A two-part practical, hands-on course on the R programming language, plotting, and accessing public databases | 138 |
| From benchside to Rstudio: Part2 |  |  | 103 |
| Introduction to Python for Biological Data Analysis: Part 1 | Hands-on | A two-part introductory workshop on Python designed for those with no prior programming experience. | 182 |
| Introduction to Python for Biological Data Analysis: Part 2 |  |  | 147 |
| Next Gen Sequencing for Biologists Series: RNA-seq Standard Analysis | Hands-on | Basics on RNA-seq data analysis. From Fastq files to differential analysis. | 122 |
| Introduction to RNA-seq Analysis on the Yale Clusters | Hands-on | Covered how to analyze RNA-sequencing data on the Yale clusters. | 59 |
| Post-GWAS analysis through integration of genome functional annotations | Demo | This session presented a statistical framework to produce high-resolution, single tissue annotations through integration of diverse epigenomic and transcriptomic data. | 34 |
| TCGA RNA-Seq Data Download and Analysis, All on Your Laptop | Hands-on | This session covered: navigation of the new The Cancer Genome Atlas data-portal and introduction of its analysis pipelines. | 71 |
| Clustering under the hood | Presentation | This introductory session focused on important statistical frameworks behind clustering of single-cell data (e.g. t-SNE, UMAP) | 70 |
| Understanding tSNE & UMAP: Comparing clustering techniques for single-cell RNA-seq | Presentation | This session shed light on tSNE and UMAP when applied on the same datasets and discussed the inner workings and some of the best practices. | 46 |
| Latex Primer | Hands-on | This session covered: use a template to produce professional looking content & eliminate formatting issues, templates provided by journals, and use the online platform "Overleaf" to produce documents on LaTeX | 20* |
|  |  | **Total** | **1062** |

(\* Capped class, no wait list enabled)

**Figure 1.**
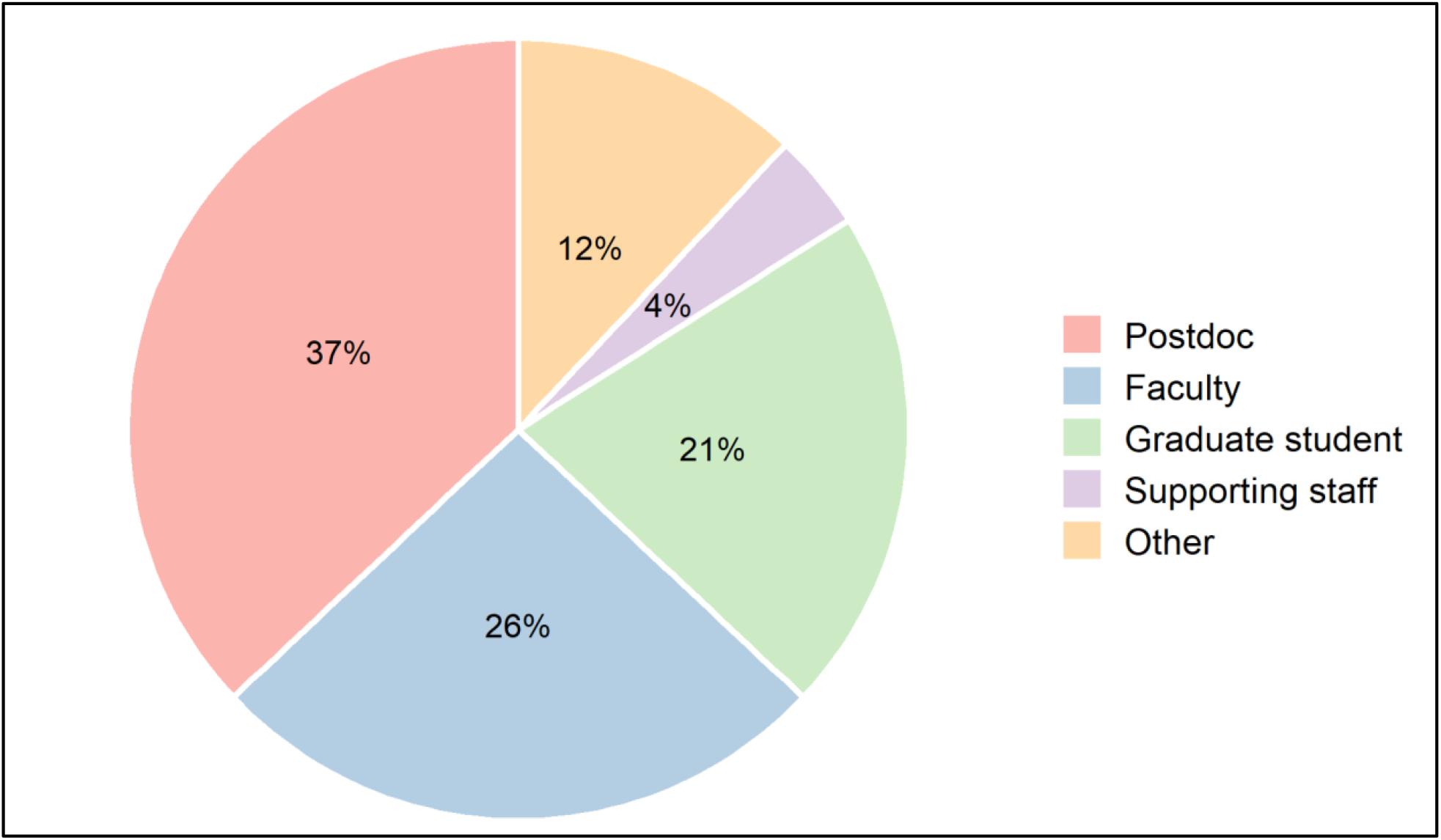
Proportion/percentage of registrants by their positions. *“Supporting staff” category includes laboratory technician, administrative assistant, and manager*. *“Other” category includes clinical research affiliate, bioinformatics technologist, hospital resident, investment analyst, programmer analyst, senior analyst, visiting research scientist, and visiting scholar*

**Figure 2.**
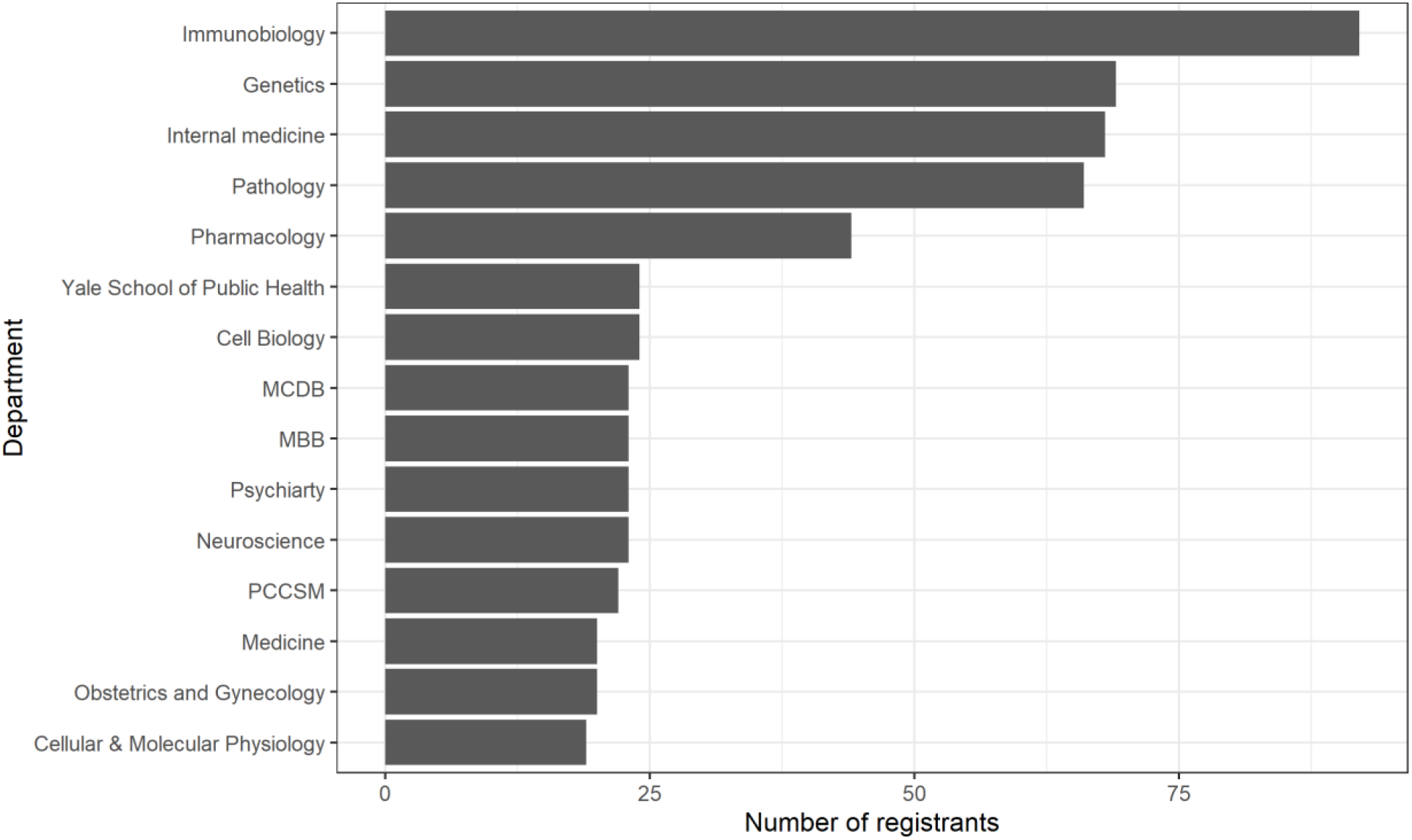
Number of registrants for most highly represented departments *MCDB: molecular, cellular, and developmental biology; MBB: molecular biophysics and biochemistry; PCCSM: pulmonary, critical care, and sleep medicine*.

## Results and Discussion

Training is one of the main aspects of the CWML Bioinformatics Support Program (https://library.medicine.yale.edu/bioinformatics). Relevant methodologies and tools are the focus of our regular hands-on workshops, demonstrations, and webinars. Training is provided by library staff, vendors, developers, and internal/external academic researchers. The rapid development and complexity of bioinformatics fields makes it impossible for any one individual to cover the wide variety of bioinformatics methods and tools. This, in addition to the unmet instructional gap revealed by a previous study (Garcia-Milian et al., 2018), made P2P teaching one of the strategies to consider.

Seven instructors responded to the call to share their bioinformatics knowledge as peer teachers. A total of 1062 individuals registered for the P2P Teaching Series. These workshops were offered between August 2018 and March 2020, and consisted of 12 unique sessions including hands-on workshops, and demonstrations. Some of these were repeated (e.g., From Benchside to Rstudio and Introduction to Python for Data Analysis) because of the high demand (Table 1).

Thirty-seven percent of registrants were postdoctoral trainees (Figure 1), even though they represent only the 17% of the YSM population (Office of Communications, 2020) https://medicine.yale.edu/communications/publishing/factsandfigures/.

We previously identified that postdoctoral trainees attribute their lack of expertise in data analysis to the absence of proper training opportunities. Therefore, it is not surprising that postdoctoral trainees have a disproportionately high representation in our workshops. In keeping with our previous findings, our most popular P2P teaching sessions were R, Python, and RNA-sequencing data analysis, topics that postdoctoral trainees had identified they need training in (Garcia-Milian et al., 2018).

Immunobiology was the most highly represented department followed by Genetics, Internal Medicine, Pathology, and Pharmacology. The rest of the top represented departments can be seen in Figure 2.

### Peer teachers’ Perspectives

Delivering good training goes beyond traditional lectures and resource-centric demos. Cooperative learning, interactivity, and problem-solving exercises can substantially enhance training quality and learning outcomes (Via et al., 2013). Since the goal of bioinformatics training is to help trainees develop long-term skills, P2P teaching is an effective model for this multi-disciplinary field.

We used the peer teachers’ (NR, DT, WLC, ATC, RO) responses to a set of open-ended questions (Byczkowska-Owczarek, 2014) to gain more insight into their personal experiences. Responses were analyzed by authors RGM, CM, and NR for commonalities and unique topics.

### Incentives to participate

Volunteering as a peer teacher in our P2P teaching program had two obvious incentives: (1) the peer teachers could upgrade their CV/resume with their teaching experience, and (2) they received a signed letter from the CWML Director, expressing gratitude for their service to the library. However, through our questionnaire we found there were two additional factors that motivated our peer teachers to volunteer in our program: the opportunity to efficiently share their expertise, and their drive to improve their own knowledge skills.

The opportunity to more efficiently share expertise manifested in two ways. For example, instead of spending valuable time in unsustainable one-on-one interactions, the teachers could deliver their knowledge to a larger group of people, which also afforded them the chance to practice communicating complicated ideas to a non-expert audience. Despite the documented necessity of proper communication by professional scientists, development of communication skills in science remains a well-known gap in doctoral- and postdoctoral-level training (Kuehne et al. 2014) (McCartney et al. 2018). The demands of doing research (such as limited time to seek other opportunities) often make it challenging to acquire proficiencies in teaching and communication.

Our findings suggest that participating as teachers in P2P teaching programs can help trainee scientists acquire vital communication skills. In the context of providing bioinformatics training, all our teachers were challenged to develop strategies that would make a complex and niche topic understandable to an audience of non-uniform expertise in bioinformatics.

As mentioned earlier, skills in bioinformatics analysis are not common nor well-distributed within institutions. Therefore, it was unsurprising to see that instructors WLC and DT, who had taught themselves programming skills were in most popular not only to their peers, but to faculty as well. One instructor detailed the frustrations and inefficiencies of the one-on-one instruction model: “*I found myself teaching and re-teaching the same methods and material*” only to find themselves “*approached by even more collaborators seeking help with their data analysis, wanting to learn bioinformatics techniques for next generation sequencing analysis*.” The proposition to use peer teaching as a solution to this problem was accepted *“enthusiastically*”.

Another incentive that emerged as a common theme among the teachers’ responses was further development of their own skills and knowledge. Multiple instructors reported that an upcoming session was “*the impetus to learn the topic thoroughly*”, so they would be able to successfully teach it. Other instructors noted that participating in the program was the perfect opportunity to “*brush up on basic programming skills that [they] might have forgotten*” and practice debugging code on the spot. In addition, one instructor mentioned that they wanted to further develop their teaching skills, which were not possible to develop solely in the lab, while another one realized the necessity for wet lab scientists to learn methods as technology progresses to “*be of service to our local scientific community*”.

### Challenges

Two primary challenges emerged in the responses from our peer teachers: time constraints due to lab workload, and classroom management. Teachers unanimously identified time constraints as the biggest barrier to more frequent peer teaching participation. Both P2P teaching program attendees and teachers wanted longer and more detailed sessions, a format that we could not accommodate due to the teachers’ research responsibilities. It is known that doctoral students often cite time constraints as the primary obstacle to participating in professional opportunities (e.g., outreach, communication skills) beyond learning how to conduct high-quality research (Andrews et at., 2005). Departments generally do not recognize that peer teaching is a skill-building opportunity that should be encouraged to benefit not only the teacher, but also their own community of researchers. As one of the teachers commented: *“there are also benefits to the lab as [a] whole, including better informing their collaborators and recruiting future lab members*.*”* The other challenge mentioned by teachers was related to classroom management. Classroom management is a broader, umbrella term describing a teacher’s efforts to oversee a multitude of activities in the classroom, including learning, social interactions, and student behavior (Martin et al., 2016). The student to teacher ratio was a particularly important consideration for us, since paying attention to individual attendees’ questions and technical issues is difficult to scale without multiple teaching assistants. As one of the teachers said, “*the main thing that would’ve made the workshop better is having more teaching assistants roaming the audience to help with debugging code*”. Furthermore, ensuring attendees’ prior knowledge of coding ahead of the class presented a hurdle — despite posting workshop prerequisites — sometimes there was a knowledge gap around basic theoretical concepts. Another aspect of classroom management was centered on logistics, such as access to servers and core software in real time: “*logistics do not always work out as expected. This forced me to learn to be fluid in designing classes, disseminating the information, and always having a back-up methodology if everything doesn’t work out as expected*.”

### Benefits

Despite the reported challenges, our teachers acknowledge that participating in the P2P teaching program was a beneficial experience. The most-noted benefit was increased confidence in public speaking, teaching, and communicating complicated concepts. One teacher even mentioned that they feel confident now to lecture on relatively new concepts rather than only on their established expertise. Another theme that emerged was increased flexibility in teaching methods. Instructors indicated that they learned how to tactfully move individual issues to the end of the session to get the group through the planned content. Similarly, the importance of having backup plans ready if technical issues occur was underscored. Our experience and findings clearly highlight that P2P teaching in library instruction has the potential to be a key tool of professional development for the peer teachers.

## Conclusions

Peer teaching is an alternative to consider for meeting the bioinformatics training needs of biomedical researchers in an academic institution. Besides the benefits of the shared knowledge for the community, it also benefits those who participate as teachers. By improving communication and pedagogical skills, these sessions become a great opportunity for their professional development.

## Author Contributions

RG-M introduced the concept and practice of peer teaching at the CWML; NR and CM contributed equally to the preparation of the manuscript, RG-M also contributed to writing; DT, WLC, ATC, and RO, all were peer teachers who participated in data collection and manuscript preparation and revision.

## Acknowledgements

We are very grateful to CWML’s Assistant Director of Research and Education Services, Judy Spak, MLS for taking the time to critically review our paper and offer her valuable advice. Also, this work could not have been possible without significant data collection by Lei Wang, MSI, CWML’s Assistant Director of Technology and Innovation services.

## Notes

### Competing Interest Statement

The authors have declared no competing interest.

